# Expression of a STING Gain-of-function Mutation in Endothelial Cells Initiates Lymphocytic Infiltration of the Lungs

**DOI:** 10.1101/2023.07.27.550897

**Authors:** Kevin MingJie Gao, Kristy Chiang, Filiz T Korkmaz, Harish Palleti Janardhan, Chinmay M. Trivedi, Lee J. Quinton, Sebastien Gingras, Katherine A. Fitzgerald, Ann Marshak-Rothstein

**Author notes:** Equal contributions (Katherine A. Fitzgerald, Ann Marshak-Rothstein). Author Contributions KMG, AMR, and KAF were involved in conceiving the project and planning the manuscript. KMG designed experiments and performed analyses. KMG, KC, and FTK performed experiments. HPJ and CMT provided guidance on investigation of endothelial cells in vivo. SG designed and generated the conditional knock-in SAVI mouse models. KMG, LJQ, AMR, and KAF interpreted and discussed the results and all authors commented on this manuscript.

## Abstract

Patients afflicted with STING gain-of-function mutations frequently present with debilitating interstitial lung disease (**ILD**) that is recapitulated in mice expressing the STING^V154M^ mutation (**VM**). Prior radiation chimera studies revealed an unexpected and critical role for non-hematopoietic cells in the initiation of ILD. To identify STING-expressing non-hematopoietic cell types relevant to ILD, we generated a conditional knock-in (**CKI**) model in which expression of the VM allele was directed to hematopoietic cells, fibroblasts, epithelial cells, or endothelial cells. Only endothelial cell-targeted expression of the mutant allele resulted in the recruitment of immune cells to the lung and the formation of bronchus-associated lymphoid tissue, as seen in the parental VM strain. These findings reveal the importance of endothelial cells as instigators of STING-driven lung disease and suggest that therapeutic targeting of STING inhibitors to endothelial cells could potentially mitigate inflammation in the lungs of SAVI patients or patients afflicted with other ILD-related disorders.

**Summary:** Patients with STING gain-of-function (GOF) mutations develop life-threatening lung autoinflammation. In this study, Gao et al. utilize a mouse model of conditional STING GOF to demonstrate a role for endothelial STING GOF in initiating immune cell recruitment into lung tissues of SAVI mice.

## Introduction

The cGAS-STING pathway is a cytosolic dsDNA sensing pathway that initiates protective inflammatory responses against viruses, bacteria, and cancer^1^. However, activation of the cGAS-STING pathway has been increasingly tied to a range of pathologic inflammatory processes across multiple organs including the gut^2^, lung^3, 4^, and brain^5–7^, pointing to the need to better understand how STING activation in specific cell types contributes to tissue pathology. One notion to explain the tissue-specific heterogeneity of cGAS-STING driven immune responses is that non-hematopoietic cell types within tissues may direct distinct inflammatory outcomes. In support of this, models of vaccination^8^ and cancer immunotherapy^9, 10^ indicate a prominent role for cGAS-STING in non-hematopoietic cells in promoting B cell and cytotoxic CD8 T cells responses, respectively. However, much remains unknown about how and which non-hematopoietic cell types mediate these cGAS-STING driven effects.

Patients with constitutively active mutations in STING develop a severe autoinflammatory disease known as STING-associated vasculopathy with onset in infancy (**SAVI**), which presents as a highly penetrant and often lethal interstitial lung disease (ILD) strongly associated with the development of cutaneous vascular endothelial pathology^11, 12^. Mice heterozygous for SAVI STING mutations similarly develop severe ILD characterized by peri-broncho-vascular bronchus-associated lymphoid tissue (BALT) formation^13, 14^. We and others have shown that ILD in SAVI mice requires lymphocytes, particularly IFNγ-producing T cells, which promote ILD and mortality^15, 16^. However, STING GOF in hematopoietic cells is not required to initiate ILD^15, 17^. Instead, we found that STING GOF in yet unidentified non-hematopoietic radioresistant cells was sufficient to initiate the activation and recruitment of lymphocytes to the lung^15^.

Presumably, the non-hematopoietic cells that drive ILD are cell types that normally express STING even in the absence of the VM mutation. Within non-hematopoietic cells of the human lung, STING expression is prominently observed in endothelial cells, epithelium, and fibroblasts^18, 19^. Importantly, activation of the cGAS-STING pathway in these non-hematopoietic cells has been associated with various infectious and sterile inflammatory pathologies. In severe SARS-COV2, activation of cGAS-STING in epithelial and endothelial cells by mitochondrial DNA promotes macrophage driven immune pathology^20, 21^ and in TNFα-induced peritonitis, STING expressing endothelial cells promote the recruitment of T lymphocytes by enhancing trans-endothelial migration^22^. Moreover, STING activation in fibroblasts has been implicated in lung fibrosis^23^. These studies identify numerous mechanisms through which STING activation in non-hematopoietic cells promotes immunopathology.

Given the evidence of prominent vasculopathy in both SAVI patients and mice, we sought to test the hypothesis that STING GOF in endothelial cells would be sufficient to initiate ILD pathology. For this purpose, we developed a novel mouse model wherein the expression of the SAVI STING V154M mutation is dependent on the activity of Cre-recombinase but remains under the control of the endogenous STING promoter (Conditional Knock-in SAVI; **CKI**). This allowed us to target the expression of the SAVI STING mutation via tissue specific Cre-recombinases. We find that targeted expression of SAVI STING in endothelial cells is sufficient to initiate the formation of lung immune aggregates seen in SAVI ILD. However, ubiquitous targeting of the SAVI mutation further exacerbated inflammation and organization of immune infiltrates within lung tissues. These findings underscore the unique and critical role of STING in endothelial cells in the initiation of ILD through the recruitment of immune cells, and demonstrates that STING GOF in additional cell types further amplifies SAVI ILD.

## Results

### Ubiquitous targeting in a mouse model of Cre-recombinase induced STING V154M expression recapitulates SAVI ILD

To assess the impact of the SAVI mutation STING V154M (VM) on specific cell types, we created a novel conditional knock-in (**CKI**) using a modified gene trap approach. This gene trap, consisting of a pair of LoxP sites flanking the adenovirus major late transcript splice acceptor^24^ followed by stop codons in each of the three reading frames, was inserted into the intron preceding exon 5 of the endogenous STING locus (Figure 1A). The VM mutation was simultaneously introduced into exon 5. Similar to other gene trap alleles, the VM allele should be null in the absence of Cre recombinase since the splice acceptor cassette creates aberrantly spliced isoforms with premature stop codons that are likely to lead to nonsense-mediated mRNA decay^25^. Heterozygous mice carrying the CKI allele still express one WT STING allele, and following Cre expression, deletion of the gene trap should restore normal splicing and expression of the VM allele. In the studies reported below, unless otherwise indicated, all mice that inherited the CKI allele also inherited and expressed one WT STING allele.

**Figure 1.**
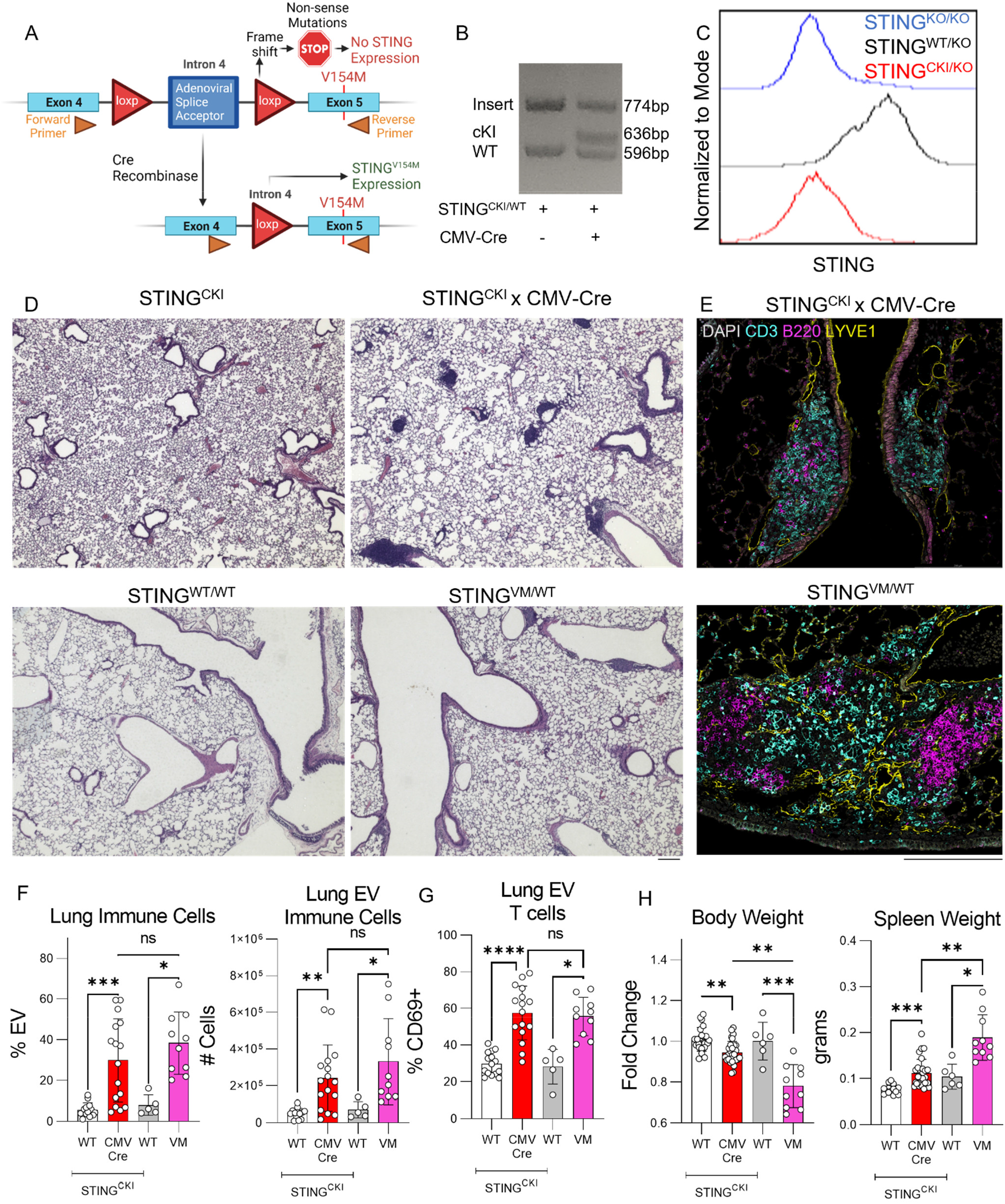
A mouse model for Cre-recombinase dependent STING V154M expression. (**A**) Diagram of the STING V154M conditional knock-in (CKI) allele demonstrating insertion of the modified gene trap cassette into the intron preceding exon 5 of the endogenous STING locus. (**B**) Tail DNA from a STING^CKI^ mouse (possessing a single copy of the CKI allele and a single copy of the STING WT allele) and a STING^CKI/WT^ x CMV-Cre mouse was PCR-amplified using primers spanning a region from exon 4 to exon 5 (indicated in diagram A). The STING WT gives a 596bp fragment, the STING CKI allele gives a 774bp fragment, and upon deletion of the gene trap from the CKI allele, a 636bp fragment is generated. (**C**) STING staining intensity in CD45+ immune cells from the blood of mice possessing either no copies of STING (STING KO/KO), a single copy of the CKI allele (STING CKI/KO), or a single copy of WT STING (STING WT/KO) as assessed by flow cytometry (D-H) 8-week-old age-, sex-, and littermate-matched STING CKI (n=13-23, white) and STING CKI x CMV-Cre mice (n=16-29, red) and 12-week-old age-, sex-, and littermate-matched WT (n=6, gray) and STING ^VM/WT^ (n=10, pink) were evaluated by the following measures. (**D**) Representative 4x field H&E stained histology from sectioned lungs. (**E**) Immunofluorescence staining of the lung for DAPI (gray), CD3 (cyan), B220 (magenta), and LYVE-1 (yellow) on STING CKI x CMV-Cre mouse lung. 200μm bars are shown in (D-E) for scale. (**F**) Percentage of lung EV immune cells from total Live CD45+ lung immune cells, and total cell counts of Live CD45+CD45IV-lung extravascular (EV) immune cells. (**G**) Percentage of CD69+ cells from lung EV CD3+ T cells. (**H**) Body weight from mice normalized as the fold change compared to the mean body weight of sex-matched STING CKI and WT controls, and spleen weight Nonparametric Mann-Whitney U-tests were used for pair-wise comparisons to determine statistical significance in F-H (ns: not significant p>0.05, *p<0.05, **p<0.01, ***p<0.001, ****p<0.0001)

As predicted, PCR amplification of CKI tail DNA using primers flanking the gene trap cassette insertion site (Figure 1A) identified an amplification product consistent with the presence of the insert (Figure 1B). Intracytoplasmic flow cytometry further confirmed the loss of STING expression in mice that only expressed the CKI allele (STING^CKI/KO^) (Figure 1C). To demonstrate that mice inheriting the CKI allele develop lung disease, we crossed CKI mice to CMV-Cre to drive ubiquitous Cre-recombinase expression. Excision of the gene trap cassette in CKI x CMV-Cre mice resulted in the detection of a new 636bp amplification product, as predicted by the activity of Cre-recombinase (Figure 1B). Furthermore, we observed that CKI x CMV-Cre mice developed ILD with prominent peri-broncho-vascular immune infiltrates (Figure 1D) rich in CD3+ and B220+ lymphocytes (Figure 1E), not observed in CMV-Cre^-/-^ x CKI control mice and comparable to the original VM mice. To quantify the number of infiltrating cells, mice were injected i.v. with a fluorophore-conjugated CD45 antibody, 3 minutes prior to euthanasia. The unstained extravascular (**EV**) CD45+ cells could then be distinguished from stained intravascular (**IV**) CD45+ cells^26^. These studies showed that CKI x CMV-Cre mice had a significant increase in the number of EV immune cells in the lung, similar to VM mice (Figure 1F). Additionally, lung EV T cells within CKI x CMV-Cre mice showed significant upregulation of the activation marker CD69, consistent with our previous studies in unmanipulated VM mice^27^. CKI x CMV-Cre mice also developed other features of systemic inflammation previously observed in VM mice, including low body weight and splenomegaly, although these findings were more profound in VM mice (Figure 1H). Thus, CKI mice develop a SAVI ILD phenotype when crossed to ubiquitous Cre expression.

### Lung stroma expresses STING

We previously used flow cytometry to demonstrate that STING is expressed in endothelium, epithelium, and fibroblasts^15^. To better understand the distribution of STING-expressing cells, lung sections from WT, VM, and STING-deficient mice (STING KO) were examined by immunofluorescent (IF) microscopy, using markers that distinguished between different cell types in the lung. LYVE-1 is well known as a lymphatic endothelium marker, but functions as a pan-endothelium marker in lung tissues^28, 29^. Podoplanin (PDPN) is expressed by lymphatic endothelium and type 1 alveolar epithelium in the lung^30^. A combination of LYVE-1, PDPN, and tissue morphology, identified LYVE-1+PDPN-blood vascular endothelium, LYVE-1+PDPN+ lymphatic endothelium, and LYVE-1-PDPN+ T1 alveolar epithelium. STING was expressed throughout the lung of both WT and VM SAVI mice, including blood and lymphatic endothelium as well as respiratory and conducting epithelium (Figure 2).

**Figure 2.**
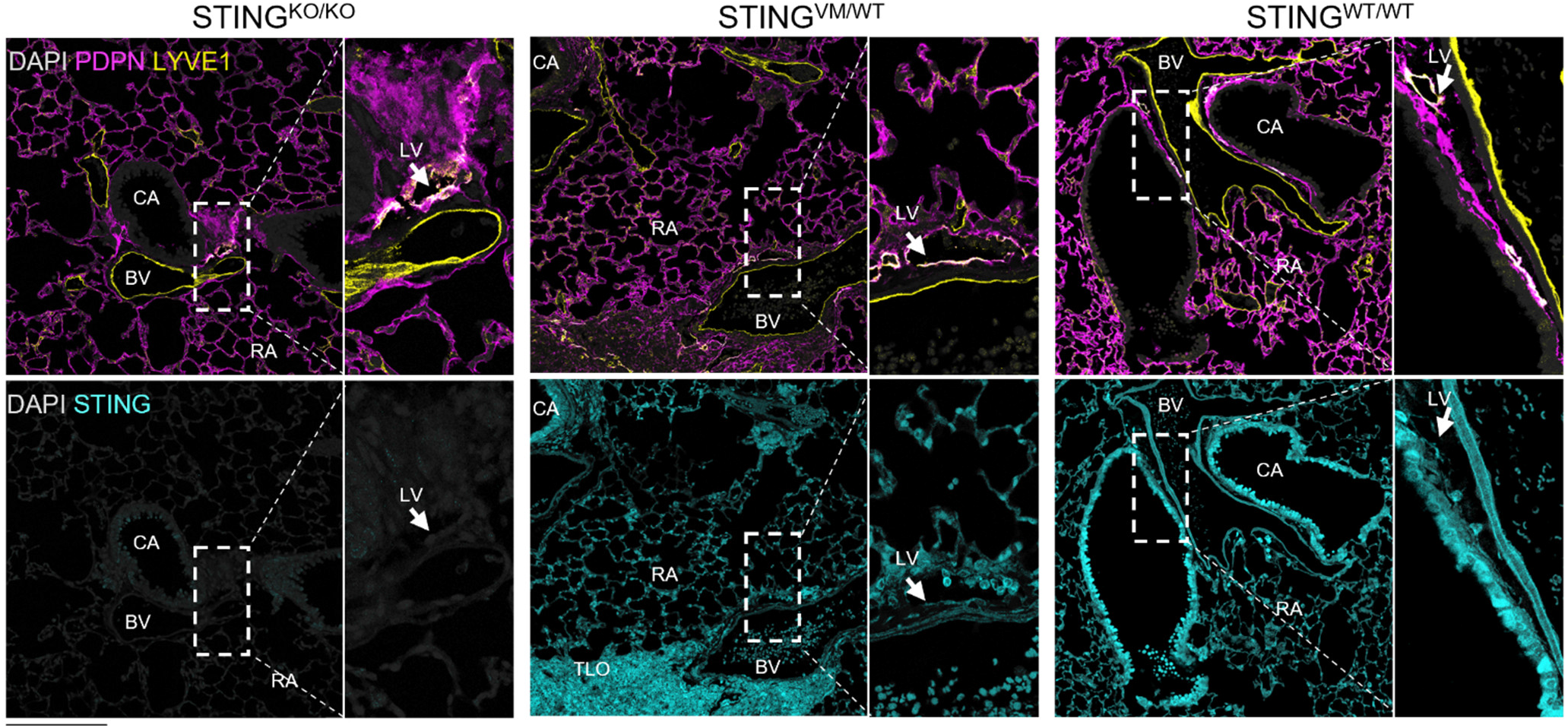
Lung stromal tissues express STING. Immunofluorescent staining of lungs from STING^KO/KO^, STING^VM/WT^, and STING^WT/WT^ mice stained for DAPI (gray), PDPN (magenta), Lyve1 (yellow), and STING (cyan). The top row shows merged DAPI, PDPN, and LYVE-1. The bottom row shows STING. A 3x magnified view is shown from the highlighted portion of each image and is shown to the right. Examples of Lyve1+PDPN-blood vessels (BV), Lyve1+PDPN+ lymphatic vessels (LV), Lyve1-PDPN-conducting airway (CA), Lyve1-PDPN+ respiratory airway (RA), and morphologically apparent tertiary lymphoid organs (TLO) are annotated by white text. A 200μm bar is shown for scale.

### Tie2-cre targeted expression of STING V154M is sufficient to initiate immune recruitment to the lung

Once we had confirmed the expression of STING across several lineages of non-hematopoietic cells in the lung, we next asked whether STING GOF in any of these cell types was sufficient to drive the SAVI ILD phenotype. We targeted lung epithelium using NKX2.1-Cre mice^31^, fibroblasts using PDGFRa-Cre mice^32, 33^, and endothelium using Tie2-Cre mice^34^. To directly measure the degree and specificity of Cre-targeting in vivo, a fluorescent Cre-reporter Rosa26-st/fl-eYFP (**YFP**) was crossed onto CKI mice to generate CKI x YFP mice. CKI x YFP mice were subsequently crossed to the tissue specific Cre lines described above. Lungs were processed by elastase digestion to optimally capture a variety of stromal and parenchymal populations within the lung^30^. As expected,CMV-Cre led to YFP reporter expression in epithelia, fibroblasts, and endothelium. Nkx2.1-Cre drove YFP reporter expression predominately in epithelium, PDGFRa-Cre in fibroblasts (as well as some epithelial cells), and Tie2-Cre in endothelial cells (Figure 3A). Thus, we confirmed that Cre-recombinase activity in Nkx2.1-Cre, PDGFRa-Cre, and Tie2-Cre targets epithelial, fibroblast, and endothelial cell types, respectively, in our CKI mice.

**Figure 3.**
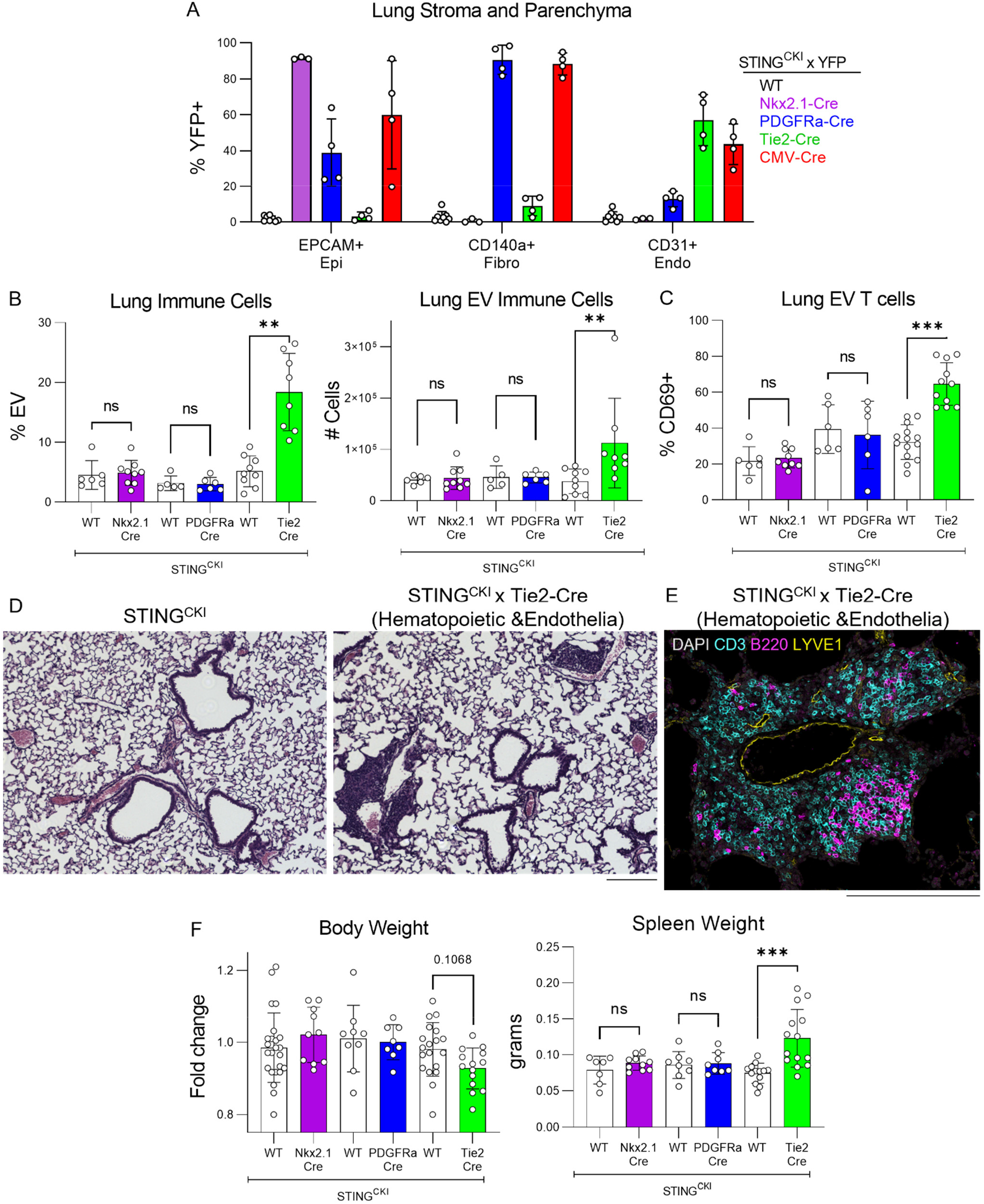
Tie2-Cre targeted expression of STING VM is sufficient to initiate immune recruitment to the lung. (**A**) Viable CD45-non-hematopoietic cells from the lungs of 12 week-old STING^CKI^ x YFP (n=8, white), STING^CKI^ x Nkx2.1-Cre x YFP (n=3, purple), STING^CKI^ x PDGFRa-Cre x YFP (n=4, blue), STING^CKI^ x Tie2-Cre x YFP (n=4, green), and STING^CKI^ x CMV-Cre x YPP (n=4, red) were evaluated for the percentage of YFP+ within EPCAM+CD31-CD140a-epithelial, CD140a+EPCAM-CD31-fibroblast, and CD31+EPCAM-CD140a-endothelial cell compartments (B-F) Three cohorts of 8-week-old mice were generated above. Cohort one contains STING^CKI^ x Nkx2.1-Cre (n=9, purple), cohort two contains STING^CKI^ x PDGFRa-Cre (n=6, blue), and cohort three contains STING^CKI^ x Tie2-Cre (n=8, green). Age-, sex-, and littermate matched control STING^CKI^ mice that do not express any additional Cre genes were generated for each cohort (n=3, n=3, n=8, respectively). These mice were evaluated by the following measures. (**B**) Percentage of lung EV immune cells within the total CD45+ lung immune cells, and total cell counts of CD45+ EV immune cells (**C**) Percentage of CD69+ lung EV T cells (**D**) Representative 10x field H&E histology from sectioned lungs of STING^CKI^ control and STING^CKI^ x Tie2-Cre mice. (**E**) Immunofluorescence staining for DAPI (gray), CD3 (cyan), B220 (magenta), and LYVE-1 (yellow) of STING^CKI^ x Tie2-Cre mouse lung. (**F**) Body weight normalized as the fold change compared to the mean body weight of sex-matched STING^CKI^ controls. Spleen weight from STING CKI x Nkx2.1-Cre, STING CKI x PDGFRa-Cre, and STING CKI x Tie2-Cre alongside their respective cohort controls. A non-parametric Kruskal-Wallis test was used for one-way ANOVA to determine statistical significance within each cohort (**p<0.01, ***p<0.001). 200μm bars are shown in (D,E) for scale.

We then asked whether we could detect lung inflammation following cell-type targeted VM expression. Targeting STING GOF to lung epithelium and fibroblasts in CKI x NKX2.1-Cre and CKI x PDGFRa-Cre mice, respectively, did not result in any increases in the number or fraction of lung EV immune infiltrates in the lungs of these mice or activation of lung EV T cells (Figure 3B,C). However, endothelial targeted STING GOF in CKI x Tie2-Cre mice was sufficient to drive an increase in EV CD45+ immune cells in the lung (Figure 3B), corresponding to the formation of histologically apparent immune aggregates (Figure 3D), and an increase in CD69 expression in lung EV T cells (Figure 3C). BALT found in these mice were enriched for T and B lymphocytes (Figure 3E), as previously observed in SAVI mice^15^. Additionally, while CKI x Tie2-Cre mice developed significant splenomegaly and showed a trend towards reduced body weight (p=0.1068), these features were not observed in the epithelial cell-targeted or fibroblast-targeted mice (Figure 3F).

### Targeted expression of STING V154M in granulocytes and T cells is insufficient to initiate ILD

The development of lung inflammation in CKI x Tie2-Cre pointed to a role for endothelial cell STING GOF in initiating SAVI ILD. However, the Tie2-promoter is also known to drive expression in hematopoietic cells^35^, and SAVI mice also exhibit aberrantly activated T cells and excessive myelopoiesis ^15, 17^. Thus, even though our previous studies, documenting the robust development of ILD in WT◊VM bone chimeras, pointed to the primary role of the VM mutation in non-hematopoietic radioresistant cells, it was important to confirm that VM expression by T cells or myeloid cells did not promote ILD.

CKI x YFP mice were crossed to Rorc-Cre (RORγ) mice to target STING GOF to T cells and innate immune lymphocytes^36^, and also crossed to LysM-Cre mice to target macrophages, monocytes, and granulocytes^37^. To confirm the specificity of T cell and monocyte/granulocyte targeting in these mice, we assessed YFP expression in splenic immune cell populations. Consistently, we found that Rorc-Cre led to prominent YFP expression in T cells; whereas, LysM-Cre led to YFP expression in neutrophils and other myeloid cells (Figure 4A).

**Figure 4.**
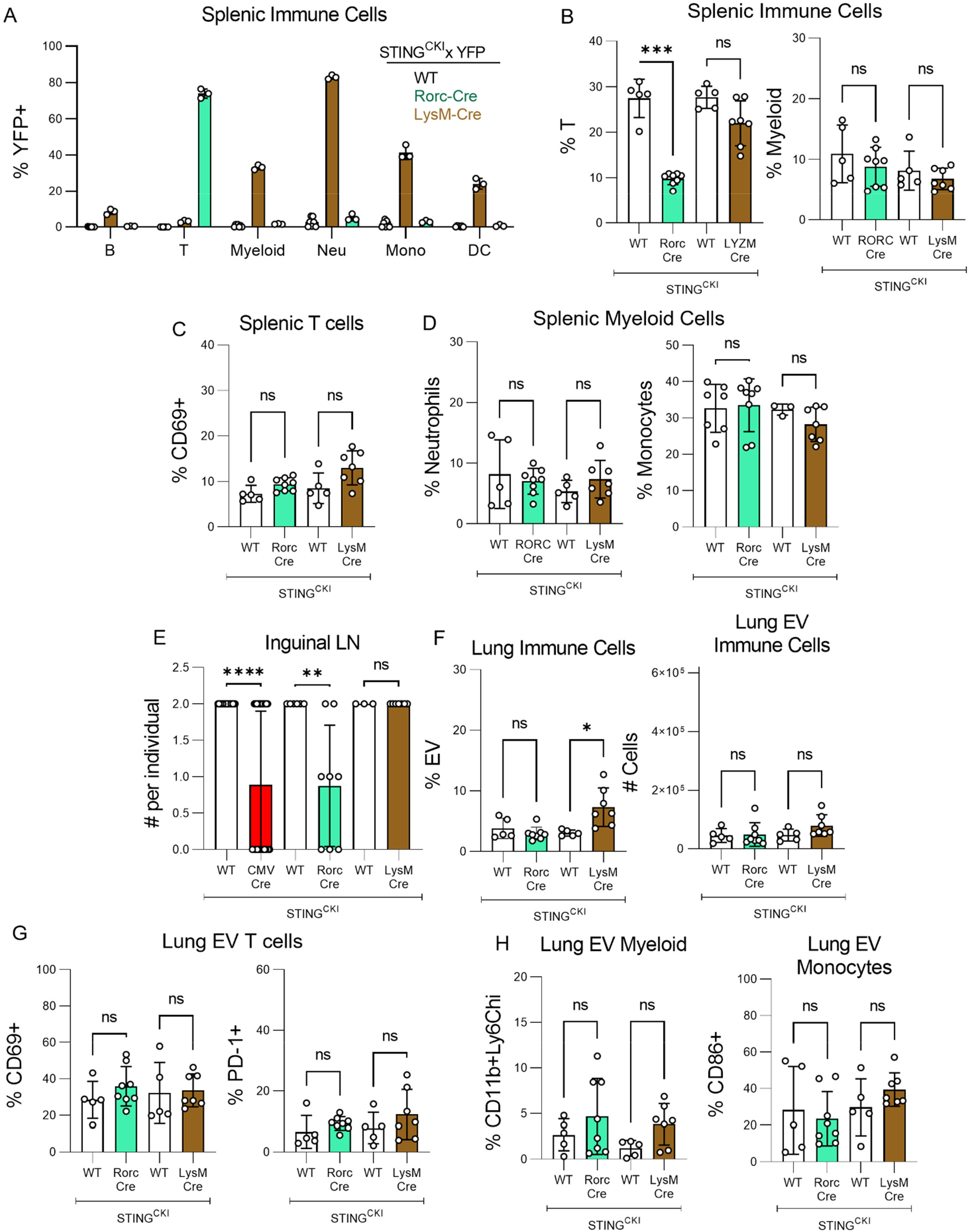
Targeted expression of STING VM in T cells and LTi induces lymphopenia and lymph node agenesis, but not ILD. (**A**) Live CD45+ splenic immune cells from 12 week-old STING^CKI^ x YFP (n=7, white), STING^CKI^ x Rorc-Cre x YFP (n=3, teal), and STING^CKI^ x LysM-Cre x YFP (n=3, brown) were evaluated for the percentage of YFP+ cells within B220+ B cell, CD3+ T cell, CD11b+ and/or CD11c+ myeloid cell, CD11b+Ly6G+ neutrophil, CD11b+Ly6G-Ly6C+ monocyte, and CD11c+MHCII+ dendritic cell subset. (B-I) Two cohorts of 8-week-old mice were generated above: STING^CKI^ x Rorc-Cre (n=8, teal) and STING^CKI^ x LysM-Cre (n=7, brown). Age-, and sex-matched control STING CKI mice that did not express any additional Cre genes were generated for each cohort (n=5, n=5, respectively). Mice were evaluated by the following measures. (**B**) Percentage of CD3+T cells and percentage of CD11b+ and/or CD11c+ myeloid cells within the total CD45+ splenocyte population. (**C**) Percentage of CD69+ within the splenic T cell compartment. (**D**) Percentage of Ly6G+ neutrophils and percentage of Ly6C+Ly6G-monocytes within the splenic myeloid subset. (**E**) Number of inguinal lymph nodes observed in comparison to a separate cohort of STING CKI (n=18, white) and STING CKI x CMV-Cre (n=27, red) mice. (**F**) Percentage of lung EV immune cells within CD45+ lung immune cells and total cell counts of EV immune cells. (**G**) Percentage of CD69+ and PD-1+ cell within EV T cell compartment. (**H**) Percentage of CD11b+Ly6C^hi^ inflammatory monocytes within lung EV myeloid compartment, and percentage of CD86+ cells within the lung EV monocyte compartment. A nonparametric Kruskal-Wallis was used for one-way ANOVA for multiple pairwise comparisons to determine statistical significance (ns: not significant p>0.05, *p<0.05,**p<0.01, ****p<0.0001).

We next determined if T cell-targeted or myeloid cell-targeted VM altered the status of immune cells in the spleens. Consistent with previous reports of lymphocyte-intrinsic STING GOF promoting T cell lymphopenia^13, 14^, we found that the number of splenic T cells was significantly reduced in CKI x Rorc-Cre mice (Figure 4B). However, the remaining T cells did not show increased expression of the activation marker CD69, as seen in T cells from unmanipulated VM mice^38^ (Figure 4C). This indicated that even though T cell-intrinsic STING GOF promotes T cell lymphopenia, it does not promote T cell activation, consistent with our previous WT◊VM radiation chimera studies showing that WT T cell activation depends on non-hematopoietic radioresistant cells in a SAVI host^15^.

Surprisingly, despite the ability of VM stem cells to give rise to an expanded myeloid compartment in VM→WT chimeras^14^, we did not find that CKI x LysM-Cre mice showed any indication of myeloid, neutrophil, or monocyte expansion in the spleen (Figure 4D). We have previously reported an increased frequency of CMP progenitors in the BM of SAVI mice^14^. Since LysM-Cre targets more mature myeloid lineage granulocytes, our finding suggests that expansion of myeloid lineage cells in SAVI likely results from STING GOF in a myeloid progenitor prior to LysM expression rather than STING GOF in mature myeloid cells.

We next asked whether T cell or myeloid-targeted VM was sufficient to induce lung inflammation. In addition to targeting T cells, Rorc-Cre also targets lymphotoxin inducer cells (LTi), a population of innate lymphoid cells (ILCs) critical for LN organogenesis^39–42^. Previous studies have shown that STING N153S SAVI mice are deficient in LTi, show impaired LN development, and have suggested that disruption of normal lymphatic function could contribute to lung inflammation in SAVI^28, 43^. We confirmed that many CKI x CMV-Cre and CKI x Rorc-Cre mice lacked inguinal LNs (Figure 4E). However, neither the CKI x Rorc-Cre nor the CKI x LysM-Cre mice exhibited increased extravascular immune infiltration of the lung, increased lung EV T cell activation markers, or signs of lung EV monocyte activation (Figure 4F-H). Thus, targeted expression of STING GOF to T cells/LTi or myeloid cells is insufficient to induce lung inflammation.

### Post-natal endothelial targeted expression of STING V154M is sufficient to initiate immune recruitment to the lung

The fact that CKI x TIE2-Cre mice, and not Rorc-Cre or LysM-Cre mice, develop features of SAVI ILD suggested that endothelial intrinsic STING GOF may indeed be responsible for initiating SAVI lung disease. However, as discussed above, constitutive Tie2-Cre is well known to have off target effects and drive recombinase activity prenatally in hematopoietic cell precursors^35^. An alternative explanation for our findings is that a combination of STING GOF in hematopoietic and endothelial cells is required to initiate ILD in CKI x Tie2-Cre mice. To address this possibility, we crossed CKI mice to a tamoxifen-inducible endothelial-targeting Cre line, Cdh5-Cre^ERT2^. Cre expression in Cdh5-Cre ^ERT2^ mice is conditional upon post-natal treatment with tamoxifen, leading to minimal off-target induction of Cre expression in cells of hematopoietic lineage while retaining endothelial-targeted Cre expression^34^. However, one important consideration of using tamoxifen-inducible Cre mice to enhance tissue-targeting specificity is that this protocol precludes targeting outcomes that occur prenatally.

To first determine whether the SAVI ILD phenotype requires an expression of STING GOF during pre-natal development, we used a ubiquitous tamoxifen-inducible Cre, CAGG-Cre^ERTM^, and bred CKI x CAGG-Cre^ERTM^ x YFP mice. These offspring were then administered three oral doses of tamoxifen postnatally on P0, P1, and P2 to induce Cre expression. When the lungs of these mice were assessed at 6-weeks of age, significant YFP expression was detected in both CD31+ endothelial and CD45+ hematopoietic cells (Figure 5A). Assessment of lung histology revealed prominent peri-broncho-vascular immune aggregates in CKI x CAGG-Cre^ERTM^ mice, but not in tamoxifen-treated CKI controls (Figure 5B). Furthermore, immunofluorescent staining of lungs from CKI x CAGG-Cre^ERTM^ mice showed well-organized BALT with distinct T and B cell zones, as seen in SAVI mice (Figure 5C). CKI x CAGG-Cre^ERTM^ also mice showed an increase in lung EV immune cells both by cell count and percentage by flow cytometry (Figure 5D). These results indicate that pre-natal expression of the VM STING mutation is not required for the development of SAVI ILD.

**Figure 5.**
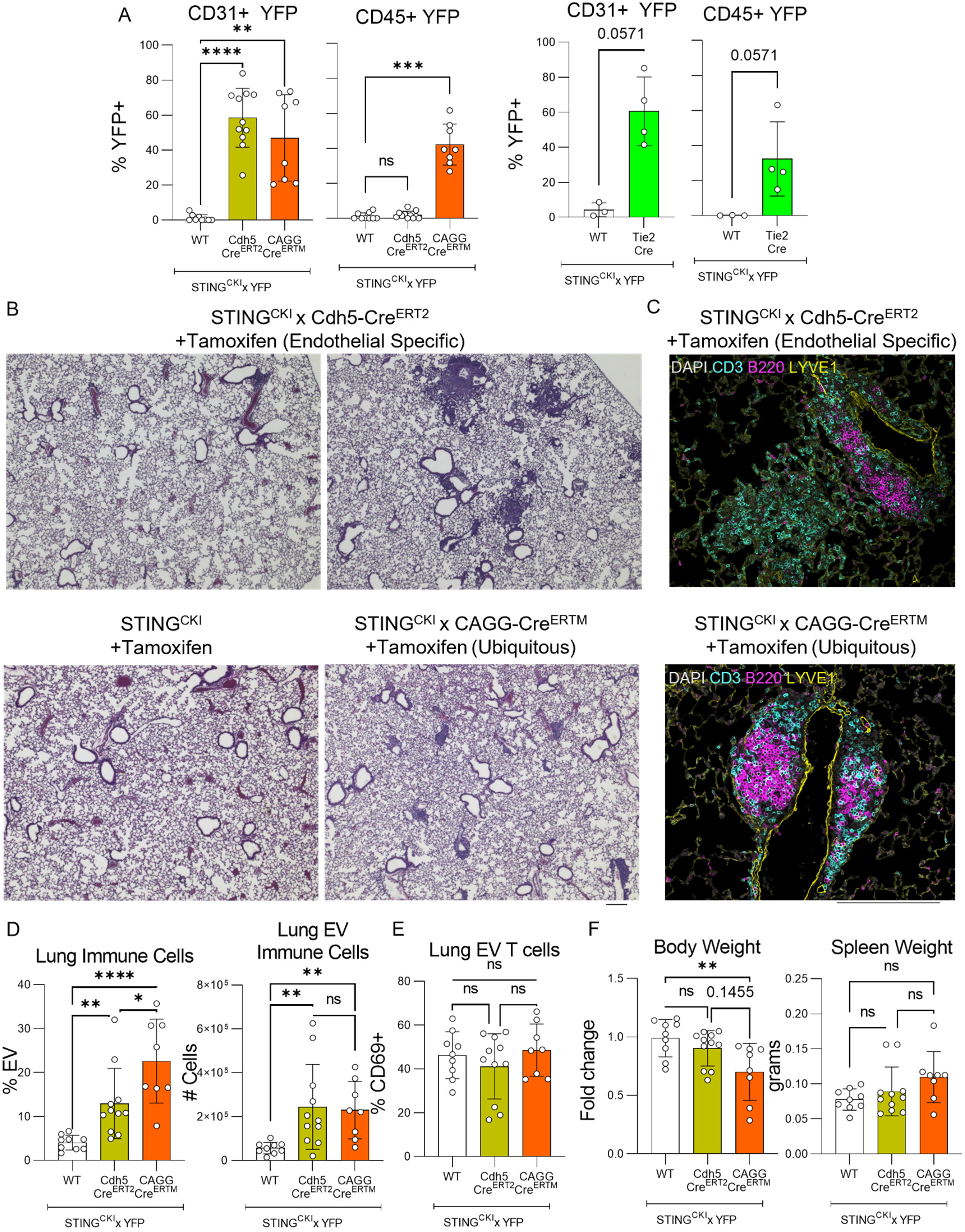
Endothelial specific expression of the STING VM mutation is sufficient to initiate immune recruitment to the lung. (**A**) The percentage of YFP+ CD31+CD45-lung endothelial cells and YFP+ CD31-CD45+ lung immune cells was assessed in two distinct cohorts: Cohort 1 (Left) consisted of sex- and littermate-controlled STING^CKI^ x YFP (n=9, white), STING^CKI^ x Cdh5-Cre^ERT2^ x YFP (n=11, yellow), and STING^CKI^ x CAGG-Cre ^ERTM^ x YFP (n=8, orange) mice, treated with tamoxifen at P0, P1, and P2 and evaluated at 5-7-weeks of age. Cohort 2 (Right) consisted of 8-week-old sex and littermate-controlled STING ^CKI^ x YFP (n=3, white) and STING ^CKI^ x Tie2-Cre x YFP (n=4, green) mice that were similarly assessed. (B-F) Cohort 1 (above) was additionally evaluated by the following criteria. (**B**) Representative 4x field H&E histology from lung sections of STING^CKI^ x Cdh5-Cre^ERT2^ and STING^CKI^ x CAGG-Cre^ERTM^ mice. Two different STING^CKI^ x Cdh5-Cre-ERT2 mice with variable presence of immune aggregates are shown: on the left is a mouse with modest immune aggregate formation, while the right is a mouse with severe immune aggregate formation. (**C**) Immunofluorescence staining for DAPI (gray), CD3 (cyan), B220 (magenta), and LYVE-1 (yellow) on STING^CKI^ x CAGG-Cre^ERTM^ and STING^CKI^ x Cdh5-Cre^ERT2^ mouse lungs. (**D**) Percentage of lung EV immune cells within total CD45+ lung immune populations, and total cell counts of total EV immune cells. (**E**) Percentage of CD69+ EV T cells. (**F**) Body weight normalized as the fold change compared to the mean body weight of sex-matched STING^CKI^ control and spleen weight Mann-Whitney U-tests were used for single pairwise comparisons, and a non-parametric Kruskal-Wallis test was used for one-way ANOVA for multiple pairwise comparisons to determine statistical significance (ns: not significant p>0.05, *p<0.05,**p<0.01, ***p<0.001, ****p<0.0001). 200μm bars are shown in (B-C) for scale.

We next compared the endothelial-targeting specificity in CKI x Tie2-Cre x YFP and CKI x CDH5-Cre-ERT2 x YFP mice. Assessment of lung cells by flow cytometry showed that CKI x Tie2-Cre x YFP mice expressed YFP in both CD31+ endothelium and CD45+ hematopoietic cells, while CKI x CDH5-Cre-ERT2 x YFP mice only expressed YFP in CD31+ cells (Figure 5A). This confirmed that Cre-activity is specifically targeted to the endothelium in CKI x x CDH5-Cre-ERT2 x YFP mice.

Importantly, histologic examination of lungs from CKI x CDH5-Cre-ERT2 x YFP mice by H&E and immunofluorescence revealed immune aggregates rich in B and T lymphocytes (Figure 5B,C). This corresponded to an increase in lung EV immune cells by absolute count and percentage compared to tamoxifen-treated CKI controls (Figure 5D). Strikingly, there was significant variability in the extent of immune cell consolidation in the lungs of CKI x CDH5-Cre ERT2 x YFP mice, with some animals showing impressively dense infiltrates (Figure 5B, top left), and others showing lightly scattered infiltrates (Figure 5B, top right). However, in both instances, there was an overall less organized appearance than the corresponding CAGG-Cre^ERTM^ mice, which showed round and condensed aggregates (Figure 5B, bottom right). This disorganized appearance was also reflected in the distribution of B and T cells observed by IF, with CKI x CDH5-Cre ERT2 mice showing poorly demarcated T and B cell zones (Figure 5C). Surprisingly, neither the CKI x CAGG-Cre or CKI x CDH5 mice showed elevated CD69 in lung EV T cell populations (Figure 5E). Lastly, CKI x CAGG-Cre-ERT2 mice showed a significant reduction in body weight and higher spleen weight (although this did not reach statistical significance), which was not observed CKI x CDH5-Cre ERT2 mice (Figure 5F). Thus, we find that targeting STING GOF to the endothelium is sufficient to initiate recruitment of immune infiltrates into lung tissue, a key feature of the SAVI ILD phenotype, but with less organized BALT compared to ubiquitous targeting and in the absence of other signs of systemic inflammation.

### Lung inflammation is enhanced by non-endothelial STING GOF

To determine whether STING GOF in additional cells beyond endothelium contributed to SAVI ILD, we compared the activation phenotypes of immune cells in the lungs of CKI x CAGG-Cre^ERTM^ x YFP and CKI x CDH5-Cre^ERT2^ x YFP mice. A hallmark of SAVI disease is the presence of activated T and B lymphocytes in the lungs of VM and WT→VM chimeric mice^15^. We found that while CKI x CAGG-Cre^ERTM^ x YFP mice showed significantly increased lung T cell expression of PD-1, this was not seen in CKI x CDH5-Cre^ERT2^ x YFP mice (Figure 6A). PD-1 is a co-inhibitory receptor which is upregulated during chronic T cell mediated inflammation^44^, thus T cell upregulation of PD-1 following ubiquitous but not endothelial specific targeting of SAVI STING expression suggests that additional factors beyond endothelial STING activation contribute to persistent activation of T cells in SAVI ILD. Moreover, PD-1 upregulation did not occur in CKI x Rorc-Cre mice, indicating that T cell extrinsic STING GOF likely accounts for this phenotype (Figure 4H). Similarly, we find that only CKI x CAGG-Cre^ERTM^ x YFP mice showed significantly increased lung B cell expression of the co-stimulatory marker CD86, again indicating that factors beyond endothelial STING GOF also promote the activation of lung B cells in SAVI (Figure 6A).

**Figure 6.**
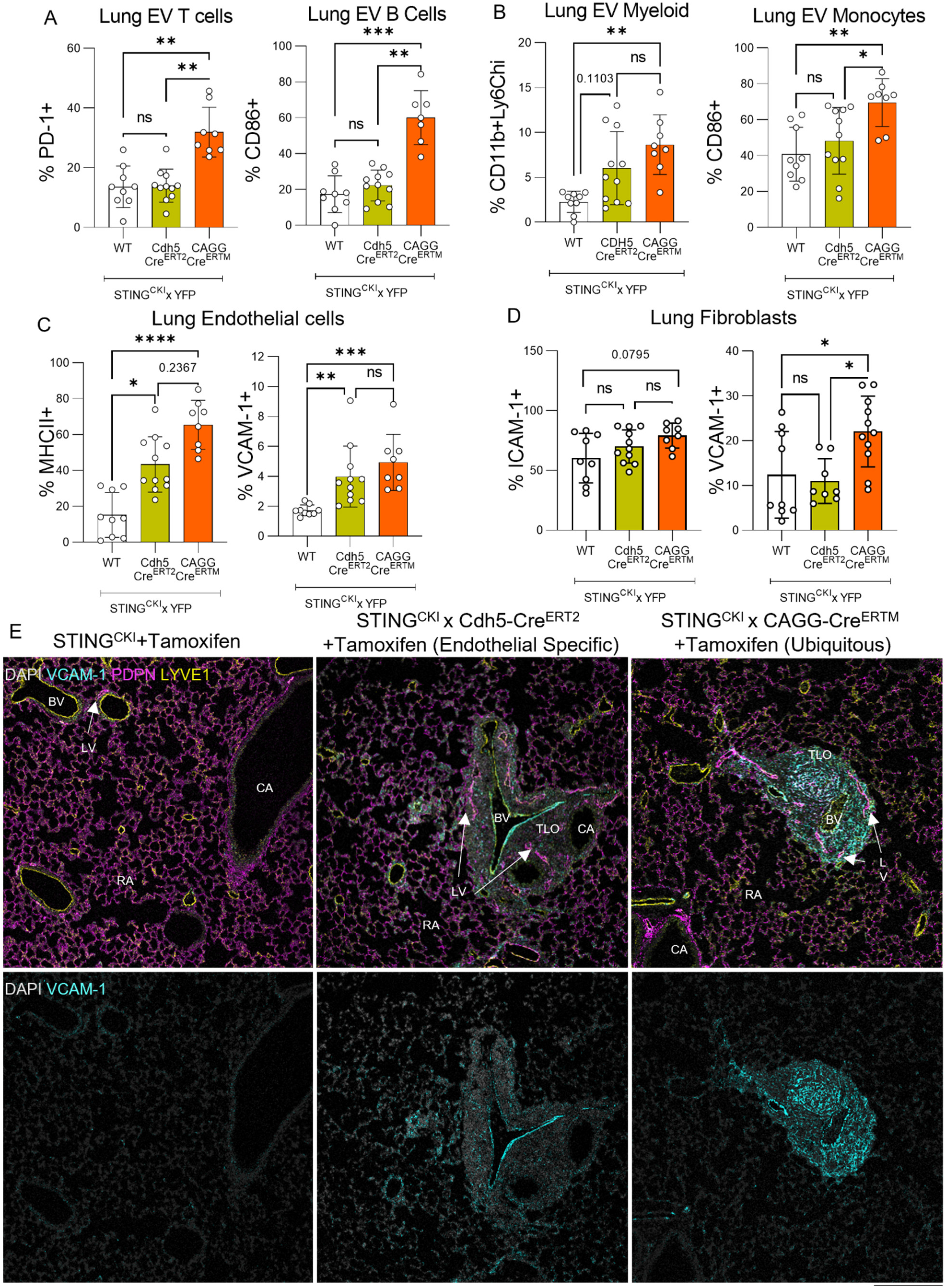
Lung inflammation is enhanced by non-endothelial expression of STING VM. A cohort of sex- and littermate-matched tamoxifen treated 6-week-old mice containing STING^CKI^ x YFP (n=9), STING^CKI^ x Cdh5-Cre ERT2 x YFP (n=11), and STING^CKI^ x CAGG-Cre ^ERTM^ x YFP (n=8) were evaluated by the following measures. (**A**) Percentage of PD-1+ EV T cells, and percentage of CD86+ CD19+ lung EV B cells. (**B**) Percentage of CD11b+ Ly6G-Ly6Chi inflammatory monocytes (IM) within the CD11b+ and/or CD11c+ EV myeloid cells, and percentage of CD86+ CD11b+ Ly6C+ lung EV monocytes. (**C**) Percentage of MHCII+ CD31+ lung endothelial cells, and percentage of VCAM1+ lung endothelial cells. (**D**) Percentage of ICAM1+ and VCAM1+ in CD31-CD140a+ lung fibroblasts. (**E**). Lung immunofluorescence All stains are shown on the top row. Only DAPI and VCAM-1 are shown on the bottom row. Nonparametric Mann-Whitney U-tests were used for single pairwise comparisons, and a nonparametric Kruskal-Wallis was used for one-way ANOVA for multiple pairwise comparisons to determine statistical significance (ns: not significant p>0.05, *p<0.05,**p<0.01, ***p<0.001, ****p<0.0001). 200μm bars are shown in (E) for scale.

Extensive infiltration of the lung by inflammatory monocytes is another distinguishing feature of SAVI ILD^15^. While lung myeloid cells in CKI x CDH5-Cre^ERT2^ x YFP mice tend to be enriched for CD11b+Ly6C hi inflammatory monocytes (p=0.1103) (Figure 6B), inflammatory monocytes are more significantly elevated in the lungs of CKI x CAGG-Cre^ERTM^ x YFP mice (Figure 6B). Moreover, only CKI x CAGG-Cre^ERTM^ x YFP mice showed upregulation of co-stimulatory ligand CD86 on lung monocytes (Figure 6B). Induction of CD86 was not observed in lung monocytes from CKI x LysM-Cre mice (Figure 4I), indicating that neither endothelial (Figure 6B) nor granulocyte (Figure 4I) intrinsic STING GOF is sufficient for exaggerated recruitment and activation of this cell type in the context of SAVI ILD. Thus, STING GOF in non-hematopoietic cells beyond those comprising the endothelium enhance monocyte recruitment and activation.

To assess the extent of endothelial cell activation per se in CKI x CDH5-Cre^ERT2^ x YFP and CKI x CAGG-Cre^ERTM^ x YFP mice, we used flow cytometry to determine the frequency of cells that had upregulated the level of MHCII expression, as a surrogate for antigen presentation capacity, along with the % of cells that had upregulated the level of the inducible adhesion molecule VCAM-1^45^, as a surrogate for leukocyte transmigration potential. Both molecules are known to be are upregulated during inflammation^46–49^. We found that expression of both MHCII and VCAM-1 were elevated in lung endothelial cells from CKI x CDH5-Cre^ERT2^ x YFP mice (Figure 6C), as well as in CKI x CAGG-Cre^ERTM^ x YFP lung endothelium (Figure 6C).

Fibroblasts also play important roles in immune responses, in particular, the organization and activation of recruited immune cells^50–53^. For example, subsets of fibroblasts known as fibroblastic reticular cells are found within lymphoid organs and express immune adhesion markers like ICAM-1 and VCAM-1 and play an important role in organizing immune aggregates^54–57^. Thus, we examined expression of these adhesion markers on lung fibroblasts as a surrogate for immune organization potential and found that ICAM-1 and VCAM-1 were both elevated in CKI x CAGG-Cre^ERTM^ x YFP but not CKI x Cdh5-Cre^ERT2^x YFP mice (Figure 6D). This observation is consistent with the more defined immune organization seen in lung immune aggregates of CKI x CAGG-Cre^ERTM^ x YFP mice (Figure 5C).

We used immunofluorescent microscopy to further compare the level of VCAM expression in these two strains. VCAM-1 staining was clearly brighter in LYVE-1+ lung endothelia of CKI x CDH5-Cre^ERT2^x YFP and CKI x CAGG-Cre^ERTM^ x YFP mice compared to CKI x YFP controls (Figure 6E). Importantly, VCAM-1 expression was limited to LYVE-1+ endothelia in CKI x CDH5-Cre^ERT2^x YFP mice, but expressed by both BALT adjacent LYVE-1+ vasculature and within LYVE-1 negative BALT stroma in the lungs of CKI xCAGG-Cre^ERTM^ x YFP mice, most likely reflecting expression by fibroblastic reticular cells (Figure 6E).

In summary, although endothelial directed STING GOF is sufficient to recruit immune cells to the lung, STING GOF in additional cell types further enhances T cell, myeloid, and fibroblast activation and contributes to lung inflammation in SAVI ILD.

## Discussion

SAVI mice recapitulate many of the features seen in human interstitial lung disease and provide an excellent model for exploring the cell-specific role of GOF STING mutations in lung inflammation. Previous radiation chimeras studies demonstrated that non-hematopoietic cells in SAVI mice were sufficient to initiate ILD^27^. Nevertheless, other reports have implicated expression of the mutant allele in T cells or innate immune cells in the development of ILD^43^. We have now utilized a novel model of conditional expression of the SAVI STING mutant V154M to demonstrate a unique role for endothelial cell expression of this STING GOF mutation in the development of SAVI ILD, specifically in the initial recruitment of immune cells to the lung. In addition, our studies also indicate that endothelial STING GOF alone does not fully account for the extent of ILD in SAVI, as ubiquitous expression of the VM mutation further amplifies the response.

The unique role of endothelial cells in SAVI ILD fits with what is known about the role of the endothelium in inflammation, as the blood endothelial barrier is tightly regulated to limit the extravasation of recruited leukocytes under homeostatic conditions^45^. However, during active inflammation, endothelial cells have been reported to upregulate contact-dependent (selectins and integrins) and contact-independent factors (cytokines) to enhance immune cell recruitment^45, 47^. T cell recognition of MHC-associated antigens also contributes to T cell recruitment by endothelial cells^58, 59^. We have now shown, by both flow cytometry and immunofluorescence, that lung endothelial cells in CKI x CDH5-Cre-ERT2 mice exhibit elevated expression of adhesion molecules like VCAM1 and antigen presentation molecules like MHCII. These observations further validate our previous finding showing gene ontology enrichment for pathways relating to immune activation (chemotaxis and antigen presentation) in VM lung endothelial cells^15^. Future work will define the functional relevance of STING activation in regulating endothelial-mediated immune recruitment processes like adhesion and trans-endothelial migration. Additionally, while blood endothelium directly contributes to the recruitment of immune cells into tissues, lymphatic endothelium plays a critical role in inflammatory resolution by promoting the drainage of immune cells from tissues back into circulation^48, 60–63^. Our future studies will also need to consider the possible contribution of lymphatic endothelium, as it is conceivable that STING GOF in lymphatic endothelium could promote inflammation by impairing the egress of immune cells out of tissues.

Previous studies from Bennion et al. using a different strategy for the conditional expression of SAVI mutant N153S found that Rorc-Cre targeting of their knock-in allele led to a loss of LTi innate lymphocytes; these mice also developed ILD. LTi plays a critical role in lymph node formation and these mice, similar to the parental SAVI mice, did not have lymph nodes^43^. One possible explanation for these findings was that normal lymphatic function is critical for resolving inflammation and maintaining tissue homeostasis^28, 62^ and that LN agenesis in SAVI mice could promote lymphatic dysfunction and the subsequent development of spontaneous lung inflammation. Although our CKI x Rorc-Cre mice also lack LN, they did not develop ILD. Importantly, expression of the VM mutant in our CKI mice is regulated by the endogenous STING locus, in contrast to the Bennion et al. model in which the mutant STING allele was constitutively expressed using the ROSA26 locus under control of the chicken actin promoter^43^. Thus, our conditional model maintains the original level and timing VM expression, which we consider a more appropriate context or delineating outcomes of STING GOF across cell types.

We were surprised to find that there was very little impact of VM expression on the composition of the myeloid compartment in CKI x LysM-Cre mice, despite neutrophilia being a prominent feature of SAVI mice^38^ and a well-known role for activation of STING in monocytes and macrophages^64^. One possibility is that the expansion of neutrophils and monocytes in SAVI mice may arise from the influence of STING GOF in myeloid progenitors, which is supported by the observation of neutrophilia in VM→WT chimeras and the increase in common myeloid progenitors within SAVI bone marrow^38^.

Our studies further indicated that endothelial STING GOF alone did not fully restore the extent of ILD seen in the original VM parental mice or in the CKI x CAGG-Cre-^ERTM^, mice where VM was expressed in all post-natal tissues. Ubiquitous post-natal expression resulted in enhanced organization of immune infiltrates and further activation of infiltrating T and myeloid cells compared to that seen with endothelial targeted expression of the VM mutant alone. We speculate that STING activation in lung epithelial cells and fibroblasts further contributes to SAVI ILD, consistent with their known functions in other settings. For example, lung epithelial cells form a critical mucosal barrier that constantly encounters foreign and self-antigens^65^. Epithelial expression of antigen presentation machinery is critical for the formation and regulation of memory T cell responses^30^. Thus, it may be that STING activation in lung epithelium promotes T cell activation. Indeed, STING agonist stimulation of the human lung epithelial cell line Calu-3 results in significant upregulation of genes involved in antigen processing and presentation^66^. Antigen presentation by fibroblasts has also been shown to facilitate memory T-cell responses^51^. Additionally, specialized fibroblasts known as fibroblastic reticular cells (FRCs) play a critical role in organizing immune responses in lymphoid organs by producing chemokines like CXCL13 and CCL19^56, 57^, and fibroblasts in the lung have also been shown to produce CXCL13 following infection^52^. STING activation in lung fibroblasts may thus play a role in the localization and persistence of the recruited immune cells to enhance inflammation. To test the combined effects of endothelial STING GOF with epithelial and/or fibroblast STING GOF in future studies, our CKI mice will be used to simultaneously target VM expression in more than one cell type by intercrossing CKI mice with mice expressing multiple tissue-specific Cre drivers.

Our observation that STING activation in endothelial cells results in immune recruitment to the lung indicates that endothelial cells initiate disease and are a prime target for therapeutic intervention. One approach for implementing this strategy would be antibody-drug conjugates, which have been utilized in the field of cancer to target chemotherapeutic agents selectively to tumors^67^. As several small molecule antagonists of STING have been recently developed (H-151^69^, SN-011^70^), it is conceivable that the conjugation of small molecule STING inhibitors to endothelial targeting antibodies could be a viable strategy to treat SAVI ILD. While inhibition of STING activation in endothelial cells would likely limit the further extravasation of immune cells, it might not shut down the ongoing lymphocyte-mediated pathogenic process. Thus, strategies to inhibit STING activation in endothelial tissue of SAVI patients would ideally be coupled to inhibitors of lymphocyte driven inflammation such as JAK-STAT inhibitors or IFNγR blockade^12, 15^.

In conclusion, our findings demonstrate a critical role for endothelial cell STING activation in mediating immune infiltration of the lung in SAVI ILD, although STING GOF in other non-hematopoietic cell types likely synergizes with endothelial STING activation to mediate fulminant disease. This finding has important implications regarding cGAS-STING sensing within endothelial cells in other pathologic settings such as infection, autoimmunity, and cancer, which can be further exploited for therapeutic benefit.

## Methods

### Mice

Rosa26-stop-eYFP (R26YFP) (Jax #006148), CMV-Cre (Jax #006054), CAGG-Cre ^ERTM^(Jax #004682), Nkx2.1-cre (Jax #008661), Tie2-cre mice (Jax #008863), Rorc-Cre (Jax #022791), and LysM-Cre (Jax #004781) were obtained from the Jackson Laboratory. PDGFRa-Cre mice (Jax #013148) were kindly provided by Dr. Jae-Hyuck Shim (UMass Chan Medical School, Worcester MA). Cdh5-Cre ERT2 mice (Taconic #13073) were kindly provided by Dr. Chinmay M. Trivedi (UMass Chan Medical School, Worcester MA). STING KO mice fully backcrossed to C57BL/6 background were kindly provided by Dr. D. Stetson (University of Washington, Seattle, WA)^77^. The parental VM mice have been described previously^14^.

The conditional V154M knock-in line (CKI) was made in by the Transgenic and Gene Targeting facility, in the Department of Immunology at the University of Pittsburgh. To generate these mice, a STOP/flox construct was inserted between exon 4 and exon 5 of the endogenous VM STING locus. Two single guide RNA were used, the SAVI-129forw target sequence (5’-gtgtggagctatgaaggctt-3’) is located in the intron upstream of the exon encoding STING-V154, the SAVI-315forw target sequence (5’-gttaaatgttgcccacgggc-3’) overlap V154. The single guide RNAs were synthesized as previously described^78^. A long single stranded oligonucleotide (synthesized by Integrated DNA Technologies) was used as donor template^79^. Briefly, fertilized embryos (C57BL/6J, The Jackson Laboratory) produced by natural mating, were microinjected, in the pronuclei, with a mixture of 0.33 µM EnGen Cas9 protein (New England Biolabs, Cat.No. M0646T), two Cas9 guides RNA: SAVI-129forw and SAVI-315forw (21.23 ng/µl each) and the long single stranded oligonucleotides SAVI-V154M-ssODN (10 ng/µl). The injected zygotes were cultured overnight, the next day the embryos that developed to the 2-cell stage were transferred to the oviducts of pseudopregnant CD1 female surrogates. Potential founder mice were genotyped by PCR and diagnostic restriction digestion of the PCR product, proper targeting was confirmed by Sanger sequencing. PCR with primers (previously described^14^) that span a region from Exon 4 to Exon 5 produce fragments of 595, 774 and 636 bp for the wild-type, targeted and recombined alleles, respectively. The primers sequences are 5’-GGTCCTCTATAAGTCCCTAAG-3’ and 5’-GGTCACCCTCAAATAAATAGG-3’. This strategy was adopted because of the simple and compact design using a long single stranded oligonucleotide as efficient donors for both insertion and gene replacement^80^. A similar targeting strategy has also been successfully used to conditionally express a pathogen mutation in Pacs1 (Thomas et al. 2023).

Sequence of the long single stranded oligonucleotide is as follows. LoxP site are cyan, the adenovirus major late transcript splice acceptor is in purple, exon 5 is in green; and substitutions (red) for V154M mutation, a silent mutation creating a NcoI site and silent substitution to inactivate the SAVI-315forw target sequence’s protospacer adjacent motif 5’ -gctcagtgctgagactcagactaatttaaaggttggagacctgggtgtggagctatgaaggcttctcgagataacttcgtatagcatacattatac gaagttattagggcgcagtagtccagggtttccttgatgatgtcatacttatcctgtcccttttttttccacagctcgcggttgaggacaaactcttcgc ggtctttccagtataacttcgtatagcatacattatacgaagttatgggatgatgggtttaatagcagtgctgagagcaagctggcagcaggttgg gaaagttttctgcaagagaagggctttggacatcccccttgaaagtccctcaggcccttctgctgtcttcagagcttgactccagcggaagtctct gcagtctgtgaagaaaagaagttaaatAtGgcccaTgggctTgcctggtcatactacattgggtacttgcggttgatcttaccaggtagggca cctctggatgttgatgtgt-3’

**Figure M1.**
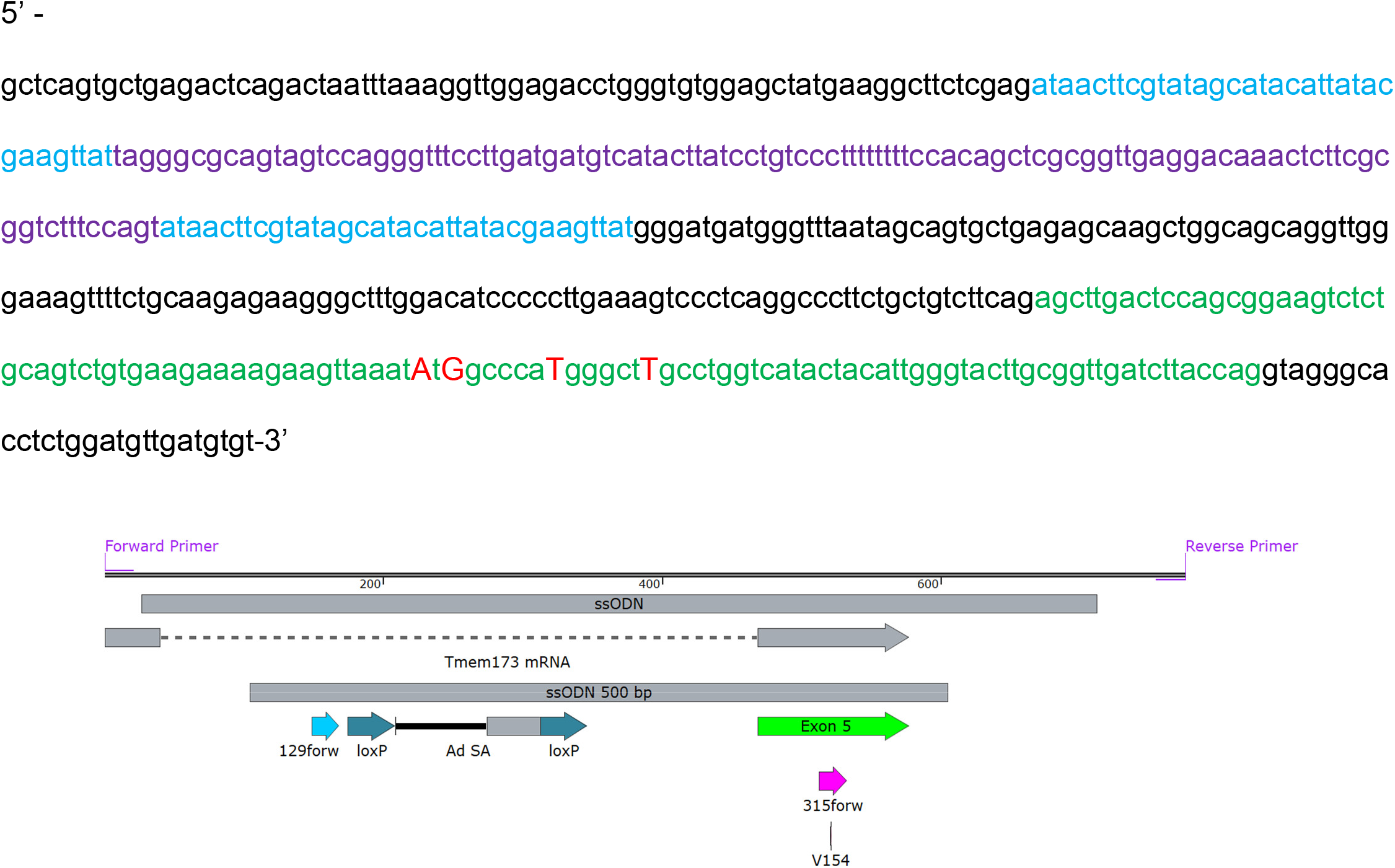
Diagram of long single stranded oligonucleotide used to generate CKI mice Homozygous founder CKI male mouse were generated and the KI allele was backcrossed to wild-type animals for 4 generations.

To prevent possible germline transmission of excised floxed cassettes in our CKI and R26eYFP mice, we took the following precautions. CKI mice were maintained as a true-breeding colony homozygous for the STING CKI allele. CKI x R26eYFP mice were generated by backcrossing CKI mice with R26eYFP mice until STING CKI homozygous R26eYFP homozygous mice were generated and maintained as true-breeding colony. We then crossed these CKI and CKI x R26eYFP true breeding mice with mice heterozygous for our various Cre-lines indicated above. The progeny from these crosses generated our experimental mice (either STING CKI/WT x Cre/WT or STING CKI/WT x R26eYFP/WT x Cre/WT) alongside littermate controls which did not carry the allele for Cre expression (either STING CKI/WT x WT/WT or STING CKI/WT x R26eYFP/WT x WT/WT). To induce the expression of Cre in mice carrying CAGG-Cre ^ERTM^ and Cdh5-Cre ERT2 alleles, we treated neonates p.o. with 2.5 ul of tamoxifen (20ug/ul) dissolved in corn oil on P0, P1, and P2.

Mice were euthanized by isoflurane followed by cervical dislocation and cardiac puncture prior to harvesting of spleen or lung tissue.. All animal experiments were conducted in accordance with the Institutional Animal Care and Use Committees at the University of Massachusetts Chan Medical School.

### Genotyping

Mice were genotyped using a combination of in-house genotyping PCRs performed on ear-clip DNA isolated from mice during weaning and custom probe-based genotyping performed by Transnetyx using real-time PCR to detect the STING V154M mutation, and the floxed STING CKI insert. Genotyping PCRs were performed on STING CKI mice using the primers indicated in Table M.1. The wildtype allele generates a 595bp fragment, the KI allele generates a 774 bp fragment, and the deleted allele generates a 636 bp fragment.

**Table M1.**
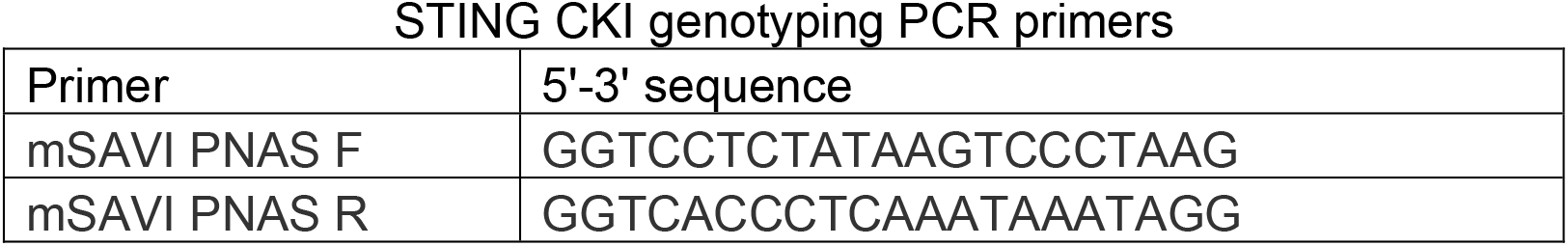
STING CKI genotyping PCR primers.

### Intravascular labelling

Circulating immune cells were identified as previously described^26^. In brief, mice were injected i.v. with 3 μg of fluorescently-labeled CD45 mAb and euthanized 3 minutes later. Lung was collected without subsequent perfusion and digested as described below.

### Lung Digestion

To assess immune cells and endothelia from the lung, the left lobe of lung was digested using a GentleMACS lung digestion kit (Miltenyi Biotec #130-095-927). In brief, lung was intratracheally inflated with 1ml of GentleMACS lung digestion kit digestion buffer. Lungs were then incubated at 37 C and dissociated using a GentleMACS Octo Dissociator with Heaters (Miltenyi Biotec #130-095-937). Cells were then filtered through a 70-micron mesh filter, spun down at 300xg for 10 minutes, and treated with RBC lysis buffer (Sigma #R7757) before subsequent assessments.

To isolate lung endothelial, epithelial, and fibroblast populations, lungs were processed and digested as previously described^30^. In brief, mice were perfused with ice-cold HBSS, and whole lung was intratracheally washed 3 times with 5 μM EDTA in DPBS. Lung was then intratracheally instilled with digestion buffer containing 4.5Units/ml of Elastase (Worthington #LS002292) and 10ug/ml Dnase I (Sigma #DN25-1g) in RPMI1640 media. Lungs were digested in a petri dish for 1 hour at 37C on a shaking platform at 200rpm, minced with a razor plate, digested for an additional 20 minutes, filtered through a 70-micron mesh filter, and pelleted by centrifugation with 300xg for 10 minutes.

### Flow Cytometry

Cells were incubated in CD16/32 (Biocell #BE0307) and stained with antibodies as documented in Tables 2 thru 4. Samples were fixed using Fluorofix buffer (Biolegend #422101). Intracellular staining was performed using a BD Biosciences Fixation/Permeabilization kit (BD Biosciences #554714) after stimulating cells for 4 hours with Brefeldin A (Biolegend #420601). Absolute cell counts were determined using counting beads (Biolegend #424902). Cells were acquired on a 5-laser Aurora (Cytek) Cytometer and analyzed with FlowJo software.

### Histology

Lungs were dissected, inflated intratracheally with 10% phosphate buffered formalin (PBF) via a flexible catheter, fixed in 10% PBF at room temperature for 48 hours, and transferred into 70% ETOH. Lungs were then paraffin embedded, sectioned, and then stained with H&E by Applied Pathology Systems (Shrewsbury, MA). Whole H&E lung slides were scanned at 4x using an EVOS FL Auto microscope or an EVOS M7000 microscope housed in the Bone Analysis Core (University of Massachusetts Chan Medical School, Worcester, MA).

### Immunofluorescence

Sections were generated from formalin fixed paraffin embedded (FFPE) lungs. For FFPE sections, 7-micron thick sections were prepared from FFPE blocks by the University of Massachusetts Chan Medical School Morphology Core. After deparaffinization using xylene, antigen retrieval was performed with 10mM Na Citrate 0.05% Tween 20 in a pressure cooker for 15 minutes. FFPE sections were then permeabilized in 0.3% TritonX-100, blocked in 10% donkey sera, incubated with primary antibody overnight at 4C, incubated with secondary antibody for 1 hour at room temperature, and then mounted with pro-long gold anti-fade with DAPI (Invitrogen #P36935) or stained with a 1ug/ml solution of DAPI (ThermoScientific #62248) and then mounted with prolong gold diamond antifade mountant (Invitrogen #P36930). Antibodies are listed in Table 5. Microscopy was captured on a Leica SP8 confocal microscope at either 10x without oil drop immersion or 63x with oil drop immersion, and then analyzed in the Leica Application Suite X.

### Statistical Analysis

Since we were unable to confirm that our data was normally distributed, we used non-parametric tests for all statistical analyses with n<15. For single comparisons, Mann-Whitney tests were performed. For one-way ANOVA, the Kruskal-Wallis test was used. For multiple comparisons testing, multiple-Mann-Whitney test was used. For Kaplan-Meier survival curve analysis, a log-rank test was used. Parametric T tests and one-way ANOVA were used when our n exceeded 15 in all groups compared.

### Antibody Panels

**Table 2.** Pan-immune flow cytometry antibody panels.

| Pan-Immune Panels |  |  |  |  |
| --- | --- | --- | --- | --- |
| Figures | Specificity | Fluorophore | Clone | Dilution |
| 1F-G<br>3C-D<br>4A-D<br>4G-I<br>5D-E<br>6A-B | IgD | PB | 11-26c.2a | 200 |
|  | PD-1 | BV421 | 29F.1A12 | 200 |
|  | Ghost | Violet510 | N/A | 500 |
|  | CD11c | BV570 | N418 | 200 |
|  | CD45.2 | BUV805 | 104 | 200 |
|  | CD45 | BV650 | 30-F11 | IV |
|  | CXCR5 | BV711 | L138D7 | 200 |
|  | CD86 | BV785 | GL-1 | 200 |
|  | Ly6G | FITC | 1A8 | 400 |
|  | CD8a | PerCP-Cy5.5 | 53-6.7 | 200 |
|  | CD11b | Spark UV387 | M1/70 | 200 |
|  | CD3 | PE | 17A2 | 200 |
|  | CXCR3 | PE-Dazzle594 | CXCR3-173 | 200 |
|  | CD69 | PE-Cy5 | H1.2F3 | 200 |
|  | CD44 | BV605 | IM7 | 200 |
|  | Ly6C | APC | HK1.4 | 300 |
|  | CD4 | SparkNIR 685 | GK1.5 | 200 |
|  | MHCII | AF700 | M5/114.15.2 | 200 |
|  | CD62L | APC-Efluor780 | MEL-14 | 200 |
|  | B220 | APC-Fire810 | RA3-6B2 | 200 |

**Table 3.** B cell flow cytometry antibody panels.

| Figures | Specificity | Fluorophore | Clone | Dilution |
| --- | --- | --- | --- | --- |
| 6A | CD138 | BV421 | 281-2 | 200 |
|  | MHCII | Pacific Blue | M5/114.15.2 | 200 |
|  | Ghost | Violet510 | N/A | 500 |
|  | Fas | BV605 | SA367H8 | 200 |
|  | CD45 | BV650 | 30-F11 | IV |
|  | CD21 | BV711 | 4E3 | 200 |
|  | CD86 | BV785 | GL-1 | 200 |
|  | GL7 | FITC | GL7 | 200 |
|  | CD11c | SparkBlue550 | N418 | 200 |
|  | CD45.2 | PerCP-Cy5.5 | 104 | 200 |
|  | IgM | PE | II/41 | 200 |
|  | CD69 | PE-Cy5 | H1.2F3 | 200 |
|  | CD23 | PE-Cy7 | B3B4 | 200 |
|  | AA4.1 | APC | AA4.1 | 200 |
|  | IgD | AF700 | 11-26c.2a | 200 |
|  | CD19 | APC-Cy7 | 6D5 | 200 |
|  | B220 | APC-Fire810 | RA3-6B2 | 200 |

**Table 4.** Non-hematopoietic flow cytometry antibody panels.

| Figures | Specificity | Fluorophore | Clone | Dilution |
| --- | --- | --- | --- | --- |
| 3A | CD8a | Spark UV 387 | 53-6.7 | 200 |
|  | MHCII | BUV563 | M5/114.15.2 | 200 |
|  | TCRb | BUV737 | H57-597 | 450 |
|  | CD11b | BUV805 | M1/70 | 200 |
|  | CD140a | BV421 | APA5 | 200 |
|  | CD4 | SparkViolet423 | RM4-5 | 200 |
|  | IgD | Pacific Blue | 11-26c.2a | 200 |
|  | Ghost | Violet 510 | N/A | 500 |
|  | Ly6C | BV570 | HK1.4 | 200 |
|  | CD45 | BV650 | 30-F11 | IV |
|  | ICAM1 | BV711 | YN1/1.7.4 | 200 |
|  | CD45.2 | BV750 | 104 | 200 |
|  | CD86 | BV785 | GL-1 | 200 |
|  | Ly6G | FITC | 1A8 | 400 |
|  | IgM | PerCP-Cy5.5 | RMM-1 | 200 |
|  | GP38 | PE | 8.1.1 | 200 |
|  | CD31 | PE-Dazzle594 | MEC13.3 | 200 |
|  | CD69 | PE-Cy5 | H1.2F3 | 200 |
|  | CD104 | PE-Cy7 | 346-11A | 200 |
|  | CD11c | PE-Fire810 | N418 | 200 |
|  | EPCAM | APC | G8.8 | 400 |
|  | CD19 | SparkNIR685 | 6D5 | 200 |
|  | MHCI | Alexa700 | AF6-88.5.3 | 200 |
|  | CD24 | APC-Cy7 | M1/69 | 200 |
|  | F4/80 | APC-Fire810 | BM8 | 100 |
| 1C<br>5A<br>6C-D | CD140a | BV421 | APA5 | 200 |
|  | CD45 | BV650 | 30-F11 | 200 |
|  | Ghost | Violet510 | N/A | 500 |
|  | Ly6C | BV570 | HK1.4 | 200 |
|  | CD45 | PB | 30-F11 | 200 |
|  | ICAM1 | BV711 | YN1/1.7.4 | 200 |
|  | CD86 | BV785 | GL-1 | 200 |
|  | ICAM2 | FITC | 3C4 (MIC2/4) | 200 |
|  | eYFP | eYFP | NA | NA |
|  | Ter119 | PerCP-Cy5.5 | TER-119 | 200 |
|  | GP38 | PE | 8.1.1 | 200 |
|  | CD31 | PE-DAZZLE<br>594 | MEC13.3 | 200 |
|  | VCAM1 | PE-Cy7 | 429(MVCAM.A) | 200 |
|  | MHCI | Alexa700 | AF6-88.5.3 | 200 |
|  | MHCII | APC_Cy7 | M5/114.15.2 | 200 |
|  | STING | Rat IgM | Clone 41 | 30 |
|  | Rat IgM | AF647 | MRM-47 | 30 |

**Table 5.** Immunofluorescence antibody panels.

| Figures | Company | Clone | Host | Isotype | Target | Conju<br>gate | Dilution |
| --- | --- | --- | --- | --- | --- | --- | --- |
| 1E<br>3E<br>5C | ABCAM | SP7/<br>ab16669 | Rabbit | IgG | CD3 |  | 150 |
|  | RND | Polyclonal/<br>AF2125 | Goat | IgG | LYVE-1 |  | 100 |
|  | BIO<br>LEGEND | RA3-6B2/<br>103204 | Rat | IgG | B220 | Biotin | 100 |
| 2 | CST | D2P2F/<br>13647 | Rabbit | IgG | STING |  | 100 |
|  | RND | Polyclonal/<br>AF2125 | Goat | IgG | LYVE-1 |  | 100 |
|  | BIO<br>LEGEND | 8.1.1/<br>127403 | Syrian<br>Hamster | IgG | PDPN | Biotin | 100 |
| 6E | ABCAM | EPR5047 | Rabbit | IgG | VCAM-1 |  | 500 |
|  | RND | Polyclonal/<br>AF2125 | Goat | IgG | LYVE-1 |  | 100 |
|  | BIO<br>LEGEND | 8.1.1/<br>127403 | Syrian<br>Hamster | IgG | PDPN | Biotin | 100 |

Secondary immunofluorescence antibodies
| Figures | Company | Clone | Host | Isotype | Target | Conjugate | Dilution |
| --- | --- | --- | --- | --- | --- | --- | --- |
| 1E<br>2<br>3E<br>5C<br>6E | Jackson | Polyclonal | Donkey | IgG | Rabbit IgG | AF488 plus | 500 |
|  | Invitrogen | Polyclonal | Donkey | IgG | Goat IgG | AF555 | 500 |
|  | Invitrogen | Polyclonal | Streptavidin | IgG | Biotin | AF647 | 500 |

## Acknowledgements

Ruth Wilson, Nicholas Mustone, Sharon Subramanian, Xhuliana Picari, and Stephanie Moses assisted in mouse breeding, genotyping and other experimental procedure. Dr. Jae Shim provided PDGFRa-Cre mice. Thank you to Dr. Kerstin Nündel, Zhaozhao Jiang, Dr. Priyadharshini Devarajan, Mingqi Dong, Kaiyuan Hao, and Jane Chuprin for constructive feedback and discussion. Chunming Bi and Zhaohui Kou of the Mouse Embryos Services facility (University of Pittsburgh School of Medicine, Department of Immunology) assisted in the development of the CKI mice. These studies were supported by: NIHAID R01 AI128358, and Lupus Research Alliance Innovation Award. KMG was supported by: National Heart, Lung, and Blood Institute (NHLBI) F30 HL154674, National Institute of General Medical Sciences (NIGMS) T32 GM107000, National Institute of Allergy and Infectious Diseases (NIHAID) T32 AI132152. CMT was supported by HL118100 and HL141377 (NHLBI). We declare the following conflicts of interest: Kate Fitzgerald is a Scientific Founder of Danger Bio, a Related Sciences company and is a member of the Scientific Advisory Board for Vesigen Therapeutics, NodThera, Janssen and Generation Bio.

## Works Cited

1. Ablasser, A. & Chen, Z. J. cGAS in action: Expanding roles in immunity and inflammation. Science (80-.). 363, eaat8657 (2019).

2. Martin, G. R., Blomquist, C. M., Henare, K. L. & Jirik, F. R. Stimulator of interferon genes (STING) activation exacerbates experimental colitis in mice. Sci. Rep. 9, 1–14 (2019).

3. Benmerzoug, S. et al. STING-dependent sensing of self-DNA drives silica-induced lung inflammation. Nat. Commun. 9, (2018).

4. Deng, Z. et al. A Defect in Thymic Tolerance Causes T Cell–Mediated Autoimmunity in a Murine Model of COPA Syndrome. J. Immunol. (2020). doi:10.4049/jimmunol.2000028

5. Chu, T. T. et al. Tonic prime-boost of STING signalling mediates Niemann–Pick disease type C. Nature (2021). doi:10.1038/s41586-021-03762-2

6. Pais, T. F. et al. Brain endothelial STING1 activation by Plasmodium-sequestered heme promotes cerebral malaria via type I IFN response. Proc. Natl. Acad. Sci. 119, e2206327119 (2022).

7. Xie, X. et al. Activation of innate immune cGAS-STING pathway contributes to Alzheimer’s pathogenesis in 5×FAD mice. *Nat*. Aging (2023). doi:10.1038/s43587-022-00337-2

8. Walker, M. M., Crute, B. W., Cambier, J. C. & Getahun, A. B Cell–Intrinsic STING Signaling Triggers Cell Activation, Synergizes with B Cell Receptor Signals, and Promotes Antibody Responses. J. Immunol. 201, 2641–2653 (2018).

9. Yang, H. et al. STING activation reprograms tumor vasculatures and synergizes with VEGFR2 blockade. J. Clin. Invest. 129, 4350–4364 (2019).

10. Yang, K. et al. Zinc cyclic di-AMP nanoparticles target and suppress tumours via endothelial STING activation and tumour-associated macrophage reinvigoration. Nat. Nanotechnol. 17, 1322–1331 (2022).

11. Liu, Y. et al. Activated STING in a Vascular and Pulmonary Syndrome. N. Engl. J. Med. 371, 507–518 (2014).

12. Frémond, M. L. et al. Overview of STING-Associated Vasculopathy with Onset in Infancy (SAVI) Among 21 Patients. J. Allergy Clin. Immunol. Pract. 9, 803–818.e11 (2021).

13. Warner, J. D. et al. STING-associated vasculopathy develops independently of IRF3 in mice. J. Exp. Med. 1–14 (2017). doi:10.1084/jem.20171351

14. Motwani, M. et al. Hierarchy of clinical manifestations in SAVI N153S and V154M mouse models. Proc. Natl. Acad. Sci. 116, 7941–7950 (2019).

15. Gao, K. M., Motwani, M., Tedder, T., Marshak-Rothstein, A. & Fitzgerald, K. A. Radioresistant cells initiate lymphocyte-dependent lung inflammation and IFNγ-dependent mortality in STING gain-of-function mice. Proc. Natl. Acad. Sci. 119, e2202327119 (2022).

16. Stinson, W. A., et al. The IFN-γ receptor promotes immune dysregulation and disease in STING gain-of-function mice. JCI insight 7, (2022).

17. Luksch, H. et al. STING-asociated lung disease in mice relies on T cells but not type I interferon. J. Allergy Clin. Immunol. (2019). doi:10.1016/j.jaci.2019.01.044

18. Digre, A. & Lindskog, C. The Human Protein Atlas—Spatial localization of the human proteome in health and disease. Protein Sci. (2021). doi:10.1002/pro.3987

19. null, null et al. The Tabula Sapiens: A multiple-organ, single-cell transcriptomic atlas of humans. Science (80-.). 376, eabl4896 (2022).

20. Di Domizio, J. et al. The cGAS-STING pathway drives type I IFN immunopathology in COVID-19. Nature (2022). doi:10.1038/s41586-022-04421-w

21. Li, H., Zhou, F. & Zhang, L. STING, a critical contributor to SARS-CoV-2 immunopathology. Signal Transduct. Target. Ther. 7, 106 (2022).

22. Anastasiou, M., Luscinskas, F. W. & Alcaide, P. Endothelial STING controls Tcell transmigration in an IFN-I dependent manner. (2021).

23. Xie, X. et al. Fluvoxamine alleviates bleomycin-induced lung fibrosis via regulating the cGAS-STING pathway. Pharmacol. Res. 187, 106577 (2023).

24. Friedrich, G. & Soriano, P. Promoter traps in embryonic stem cells: A genetic screen to identify and mutate developmental genes in mice. Genes Dev. (1991). doi:10.1101/gad.5.9.1513

25. Popp, M. W. & Maquat, L. E. Leveraging rules of nonsense-mediated mRNA decay for genome engineering and personalized medicine. Cell (2016). doi:10.1016/j.cell.2016.05.053

26. Anderson, K. G. et al. Intravascular staining for discrimination of vascular and tissue leukocytes. Nat. Protoc. 9, 209–222 (2014).

27. Gao, K. M., Motwani, M., Tedder, T., Marshak-Rothstein, A. & Fitzgerald, K. Radioresistant cells initiate lymphocyte-dependent lung in fl ammation and IFN γ -dependent mortality in STING. PNAS 119, 1–12 (2022).

28. Reed, H. O. et al. Lymphatic impairment leads to pulmonary tertiary lymphoid organ formation and alveolar damage. J. Clin. Invest. 129, 2514–2526 (2019).

29. Kalucka, J. et al. Single-Cell Transcriptome Atlas of Murine Endothelial Cells. Cell (2020). doi:10.1016/j.cell.2020.01.015

30. Shenoy, A. T. et al. Antigen presentation by lung epithelial cells directs CD4+ TRM cell function and regulates barrier immunity. Nat. Commun. (2021). doi:10.1038/s41467-021-26045-w

31. Rawlins, E. L. & Perl, A. K. The a"MAZE"ing world of lung-specific transgenic mice. Am. J. Respir. Cell Mol. Biol. 46, 269–282 (2012).

32. Swonger, J. M., Liu, J. S., Ivey, M. J. & Tallquist, M. D. Genetic tools for identifying and manipulating fibroblasts in the mouse. Differentiation (2016). doi:10.1016/j.diff.2016.05.009

33. Ntokou, A. et al. Characterization of the platelet-derived growth factor receptor-α-positive cell lineage during murine late lung development. Am. J. Physiol. - Lung Cell. Mol. Physiol. (2015). doi:10.1152/ajplung.00272.2014

34. Payne, S., Val, S. De & Neal, A. Endothelial-specific cre mouse models is your cre CREdibile? Arterioscler. Thromb. Vasc. Biol. (2018). doi:10.1161/ATVBAHA.118.309669

35. Tang, Y., Harrington, A., Yang, X., Friesel, R. E. & Liaw, L. The contribution of the Tie2+ lineage to primitive and definitive hematopoietic cells. Genesis (2010). doi:10.1002/dvg.20654

36. Eberl, G. & Littman, D. R. Thymic origin of intestinal alphabeta T cells revealed by fate mapping of RORgammat+ cells. Science (80-.). (2004).

37. Abram, C. L., Roberge, G. L., Hu, Y. & Lowell, C. A. Comparative analysis of the efficiency and specificity of myeloid-Cre deleting strains using ROSA-EYFP reporter mice. J. Immunol. Methods 408, 89–100 (2014).

38. Motwani, M. et al. Hierarchy of clinical manifestations in SAVI N153S and V154M mouse models. 1–10 (2019). doi:10.1073/pnas.1818281116

39. Vivier, E. et al. Innate Lymphoid Cells: 10 Years On. Cell 174, 1054–1066 (2018).

40. Bando, J. K. & Colonna, M. Innate lymphoid cell function in the context of adaptive immunity. Nat. Immunol. 17, 783–789 (2016).

41. Onder, L. & Ludewig, B. A Fresh View on Lymph Node Organogenesis. Trends Immunol. 39, 775–787 (2018).

42. Vivier, E., Van De Pavert, S. A., Cooper, M. D. & Belz, G. T. The evolution of innate lymphoid cells. Nat. Immunol. 17, 790–794 (2016).

43. Bennion, B. G. et al. STING Gain-of-Function Disrupts Lymph Node Organogenesis and Innate Lymphoid Cell Development in Mice. Cell Rep. 31, (2020).

44. Sharpe, A. H. & Pauken, K. E. The diverse functions of the PD1 inhibitory pathway. Nature Reviews Immunology (2018). doi:10.1038/nri.2017.108

45. Alcaide, P. Mechanisms Regulating T Cell–Endothelial Cell Interactions. Cold Spring Harb. Perspect. Med. 12, 1–10 (2022).

46. Mai, J., Virtue, A., Shen, J., Wang, H. & Yang, X. F. An evolving new paradigm: Endothelial cells - Conditional innate immune cells. J. Hematol. Oncol. 6, 1–13 (2013).

47. Amersfoort, J., Eelen, G. & Carmeliet, P. Immunomodulation by endothelial cells — partnering up with the immune system? Nat. Rev. Immunol. 1, (2022).

48. Card, C. M., Yu, S. S. & Swartz, M. A. Emerging roles of lymphatic endothelium in regulating adaptive immunity. J. Clin. Invest. 124, 943–952 (2014).

49. Santambrogio, L., Berendam, S. J. & Engelhard, V. H. The antigen processing and presentation machinery in lymphatic endothelial cells. Front. Immunol. 10, 1–9 (2019).

50. Brown, F. D. et al. Fibroblastic reticular cells enhance T cell metabolism and survival via epigenetic remodeling. Nat. Immunol. 20, 1668–1680 (2019).

51. Cupovic, J. et al. Adenovirus vector vaccination reprograms pulmonary fibroblastic niches to support protective inflating memory CD8+ T cells. Nat. Immunol. 22, 1042–1051 (2021).

52. Denton, A. E. et al. Type I interferon induces CXCL13 to support ectopic germinal center formation. J. Exp. Med. 216, 621–637 (2019).

53. Krishnamurty, A. T. et al. LRRC15+ myofibroblasts dictate the stromal setpoint to suppress tumour immunity. Nature 611, 148–154 (2022).

54. Perez-Shibayama, C., Gil-Cruz, C. & Ludewig, B. Fibroblastic reticular cells at the nexus of innate and adaptive immune responses. Immunological Reviews (2019). doi:10.1111/imr.12748

55. Onder, L., Cheng, H.-W. & Ludewig, B. Visualization and functional characterization of lymphoid organ fibroblasts*. Immunol. Rev. 306, 108–122 (2022).

56. Krishnamurty, A. T. & Turley, S. J. Lymph node stromal cells: cartographers of the immune system. Nature Immunology (2020). doi:10.1038/s41590-020-0635-3

57. Buechler, M. B. & Turley, S. J. A short field guide to fibroblast function in immunity. Seminars in Immunology (2018). doi:10.1016/j.smim.2017.11.001

58. Pober, J. S., Merola, J., Liu, R. & Manes, T. D. Antigen presentation by vascular cells. Frontiers in Immunology (2017). doi:10.3389/fimmu.2017.01907

59. Manes, T. D. & Pober, J. S. Significant differences in antigen-induced transendothelial migration of human CD8 and CD4 T effector memory cells. Arterioscler. Thromb. Vasc. Biol. (2016). doi:10.1161/ATVBAHA.116.308039

60. Schwager, S. & Detmar, M. Inflammation and lymphatic function. Front. Immunol. 10, 1–11 (2019).

61. Reed, H. O. et al. Lymphatic impairment leads to pulmonary tertiary lymphoid organ formation and alveolar damage Graphical abstract Find the latest version : Lymphatic impairment leads to pulmonary tertiary lymphoid organ formation and alveolar damage. J. Clin. Invest. 129, 2514– 2526 (2019).

62. Baluk, P. et al. Lymphatic Proliferation Ameliorates Pulmonary Fibrosis after Lung Injury. Am. J. Pathol. 190, 2355–2375 (2020).

63. Baluk, P. et al. Preferential lymphatic growth in bronchus-associated lymphoid tissue in sustained lung inflammation. Am. J. Pathol. (2014). doi:10.1016/j.ajpath.2014.01.021

64. Wu, J., Dobbs, N., Yang, K. & Yan, N. Interferon-Independent Activities of Mammalian STING Mediate Antiviral Response and Tumor Immune Evasion. Immunity 53, 115–126.e5 (2020).

65. Hewitt, R. J. & Lloyd, C. M. Regulation of immune responses by the airway epithelial cell landscape. Nat. Rev. Immunol. (2021). doi:10.1038/s41577-020-00477-9

66. Li, M. et al. Pharmacological activation of STING blocks SARS-CoV-2 infection. Sci. Immunol. 6, (2021).

67. Firer, M. A. & Gellerman, G. Targeted drug delivery for cancer therapy: The other side of antibodies. Journal of Hematology and Oncology (2012). doi:10.1186/1756-8722-5-70

68. Kiseleva, R. Y. et al. Targeting therapeutics to endothelium: are we there yet? Drug Delivery and Translational Research (2018). doi:10.1007/s13346-017-0464-6

69. Haag, S. M. et al. inhibitors. Nature (2018). doi:10.1038/s41586-018-0287-8

70. Hong, Z. et al. STING inhibitors target the cyclic dinucleotide binding pocket. Proc. Natl. Acad. Sci. U. S. A. (2021). doi:10.1073/pnas.2105465118

71. Tu, X. et al. Interruption of post-Golgi STING trafficking activates tonic interferon signaling. Nat. Commun. 2022 131 13, 1–16 (2022).

72. Kasparcova, V. et al. Design of amidobenzimidazole STING receptor agonists with systemic activity. Nature 564, 439–443 (2018).

73. Pan, B. S. et al. An orally available non-nucleotide STING agonist with antitumor activity. Science (80-.). (2020). doi:10.1126/science.aba6098

74. Woo, S. R. et al. STING-dependent cytosolic DNA sensing mediates innate immune recognition of immunogenic tumors. Immunity 41, 830–842 (2014).

75. Rodriguez, A. B. & Engelhard, V. H. Insights into tumor-associated tertiary lymphoid structures: Novel targets for antitumor immunity and cancer immunotherapy. Cancer Immunol. Res. (2020). doi:10.1158/2326-6066.CIR-20-0432

76. Kang, W. et al. Tertiary Lymphoid Structures in Cancer: The Double-Edged Sword Role in Antitumor Immunity and Potential Therapeutic Induction Strategies. Frontiers in Immunology (2021). doi:10.3389/fimmu.2021.689270

77. Ishikawa, H. & Barber, G. N. STING is an endoplasmic reticulum adaptor that facilitates innate immune signalling. Nature 455, 674–678 (2008).

78. Pelletier, S., Gingras, S. & Green, D. R. Mouse Genome Engineering via CRISPR-Cas9 for Study of Immune Function. Immunity (2015). doi:10.1016/j.immuni.2015.01.004

79. Quadros, R. M. et al. Easi-CRISPR: A robust method for one-step generation of mice carrying conditional and insertion alleles using long ssDNA donors and CRISPR ribonucleoproteins. Genome Biol. 18, 1–15 (2017).

80. Miura, H., Quadros, R. M., Gurumurthy, C. B. & Ohtsuka, M. Easi-CRISPR for creating knock-in and conditional knockout mouse models using long ssDNA donors. Nat. Protoc. 13, 195–215 (2018).

